# *In vivo* deuterated water labeling allows tumor visualization via deuterium magnetic resonance spectroscopic imaging of cholesterol

**DOI:** 10.1101/809327

**Authors:** Julian C. Assmann, Jeffrey R. Brender, Don E. Farthing, Keita Saito, Shun Kishimoto, Kathrynne A. Warrick, Natella Maglakelidze, Daniel R. Crooks, Hellmut Merkle, Ronald E. Gress, Murali C. Krishna, Nataliya P. Buxbaum

## Abstract

Water is an essential component of many biochemical reactions. Deuterated water (D_2_O) has been used to study cell kinetics, protein synthesis, and metabolism. We hypothesized that rapidly proliferating cancer cells would become preferentially labeled with deuterium due to high metabolic activity, thus allowing imaging of biosynthetically labeled metabolites within tumors *in vivo*. We initiated systemic D_2_O labeling in two established tumor xenograft models, HT-29 and MiaPaCa-2 and imaged mice by deuterium magnetic resonance spectroscopic imaging (dMRSI). After 14 days of tumor growth and 7 days of *in vivo* labeling, a clear contrast was demonstrated between the xenograft and the contralateral control limb in both models. The origin of the contrast was traced to an aliphatic peak at 1.8 ppm, which was identified by *ex vivo* NMR analysis to originate from cholesterol and cholesterol esters. Cholesterol is important for tumor cell proliferation, signaling, and malignant transformation, while current methods to monitor cholesterol synthesis and accumulation are limited. This deuterated water labeling-imaging approach could complement current cancer imaging techniques, allowing not only imaging of uptake but also synthesis of cholesterol to elucidate effects on tumor cholesterol metabolism *in vivo*.

## Introduction

Imaging is an essential tool for cancer diagnosis, staging, and surveillance. Positron emission tomography (PET) and computer tomography (CT) scans are commonly used in this setting. They provide clinically valuable information that can guide therapeutic decisions (1-3). Radioactive tracers, such as ^18^fluorodeoxyglucose (FDG), in combination with CT take advantage of unique features of cancer cells such as their highly proliferative and glycolytic nature, allowing identification of small tumors that are not detectable by anatomical imaging alone (4). Such detection can result in clinically meaningful outcomes for patients, as surgical or medical interventions can be undertaken at lower tumor burden, i.e. prior to local growth and/or metastasis.

While PET tracers can probe multiple pathways, the signal is insensitive to the exact chemical fate of the tracer and the information gained is therefore primarily limited to uptake and retention. As evidence accumulates that alterations in metabolism are critical for tumor survival and progression, alternatives to PET-CT are currently being developed that can track such alterations. This includes hyperpolarized ^13^C-MRI, a technique that uses dynamic nuclear polarization (DNP) to facilitate detection of ^13^C-labeled substrates (e.g. ^13^C-pyruvate) (5).

We hypothesized that, instead of using a labeled radiotracer, we would be able to label tumors *in vivo* biosynthetically by providing deuterated water (D_2_O) as a labeled substrate systemically and for relatively short periods to exploit the differential proliferation and metabolic rates of cancer cells and normal tissues. By definition, malignant cells exhibit rapid growth and frequent cell division. To sustain these high rates of proliferation, tumor cells must generate new cellular biomass (6,7). During biosynthesis, (deuterated) water can be used as a substrate for enzymatic reactions in multiple pathways, leading to the formation of stable carbon-deuterium bonds that are not exchangeable with hydrogen, thus allowing *in vivo* labeling of proliferating cells via simple oral administration of deuterated water (8-10). This principle has formed the basis for the decades-long use of D_2_O to study cell cycle kinetics in both patients and animal models with virtually no adverse effects reported at low to moderate concentrations, up to 30% v/v in animal studies (11-13). These studies used mass spectrometry to measure cell division rates of cells during a specified labeling period and subsequently extracted from tissues or blood for the analysis. Following extraction, deuterium incorporation into newly synthesized DNA strands was quantified to estimate how many new cells were formed during the *in vivo* labeling period.

*In vivo* imaging offers several advantages by limiting invasiveness and improving coverage compared to *ex vivo* analyses of tissue biopsies. Deuterium magnetic resonance spectroscopic imaging (dMRSI) is an emerging technique to measure metabolic changes *in vivo* by MRI. dMRSI takes advantage of the rapid relaxation of the deuterium nucleus to enable faster cycling of scans, thereby regaining some of the reduced sensitivity from its low-gamma spin (14,15). Here, we present a novel imaging approach using *in vivo* D_2_O labeling followed by dMRSI for the visualization of tumors in two xenograft mouse models. We recently used this technique to visualize target organs of graft-versus-host disease infiltrated by alloreactive T cells, which share certain features with tumor cells, i.e. rapid proliferation and glycolytic metabolism (13). Using this deuterium administration protocol with tumor-bearing mice, we quantitatively show by mass spectrometric analysis that tumor xenografts undergo a concentration-dependent enrichment of deuterium into the DNA base deoxyadenosine (dA). Furthermore, by combining deuterium labeling with dMRSI, we quantitatively showed cholesterol accumulation in two independent tumor models, demonstrating the ability of this technique to provide unique metabolic information. This relatively easy to implement, non-radioactive imaging method could provide a useful addition to the imaging arsenal for cancer research.

## Methods

### Mice

Female athymic nude (Foxn1^nu^) mice aged 10 – 15 weeks were supplied by the Frederick Cancer Research Center (Frederick, MD, USA). Mice were kept under a 12h/12h light-dark cycle with *ad libitum* access to food and water. All animal procedures were approved by the NCI Institutional Animal Use and Care Committee.

### Cells

The human colorectal adenocarcinoma cell line HT-29 was purchased from ATCC (HTB- 38) and the identity was confirmed using a panel of microsatellite markers (IDEXX Laboratories). HT-29 cells were cultured in RPMI 1640 supplemented with 10% fetal bovine serum, 100 U/ml penicillin, 100 µg/ml streptomycin and incubated at 5% CO_2_ and 37 °C. On the day of tumor injection, HT-29 cells were spun down at 1000 rpm for 5 min, resuspended in PBS and 1 × 10^6^ cultured cells were injected subcutaneously into the right proximal hind limb of the mouse as published previously (16). The contralateral hind limb of each mouse did not receive a tumor cell injection and served as an intra-individual control. Similarly, the human pancreatic cancer cell line MiaPaCa-2 (CRL-1420) was cultured as described above and 3 × 10^6^ cells were injected subcutaneously into the right hind limb.

### Deuterium labeling

The dMRSI studies were performed per the experimental schema described in Fig. 2 either 7 or 14 days after tumor cell implantation. For these imaging studies, a D_2_O level of ∼8% in total body water (TBW) was targeted. The mice initially received two bolus injections (35 ml/kg body weight) containing NaCl (0.9%, w/v) in D_2_O (100%, Cambridge Isotope Laboratories) 24 h apart, each bolus increasing the TBW enrichment by ∼4%. Thereafter, the mice were provided drinking water containing 16% (v/v) D_2_O (70% D_2_O, Cambridge Isotope Laboratories, diluted with sterile ultra-pure water, Quality Biological) until imaging was performed (8). Administration of 16% (v/v) D_2_O was necessary to maintain 8% D_2_O in TBW taking into account the 30-40% loss of D_2_O due to respiration and excretion (8). D_2_O dosing was either initiated the day prior to tumor cell injection or 6 days after as indicated in the figure legend. For the dose-escalation study, mice received 1-4 bolus injections within 7 days prior to tumor cell injection and were subsequently provided 8%, 16% and 32% D_2_O drinking water to achieve targeted concentrations of 4%, 8% and 16% D_2_O, respectively in TBW based on previous publications (8). Regular drinking water served as a control (0% D_2_O).

**Figure 1:**
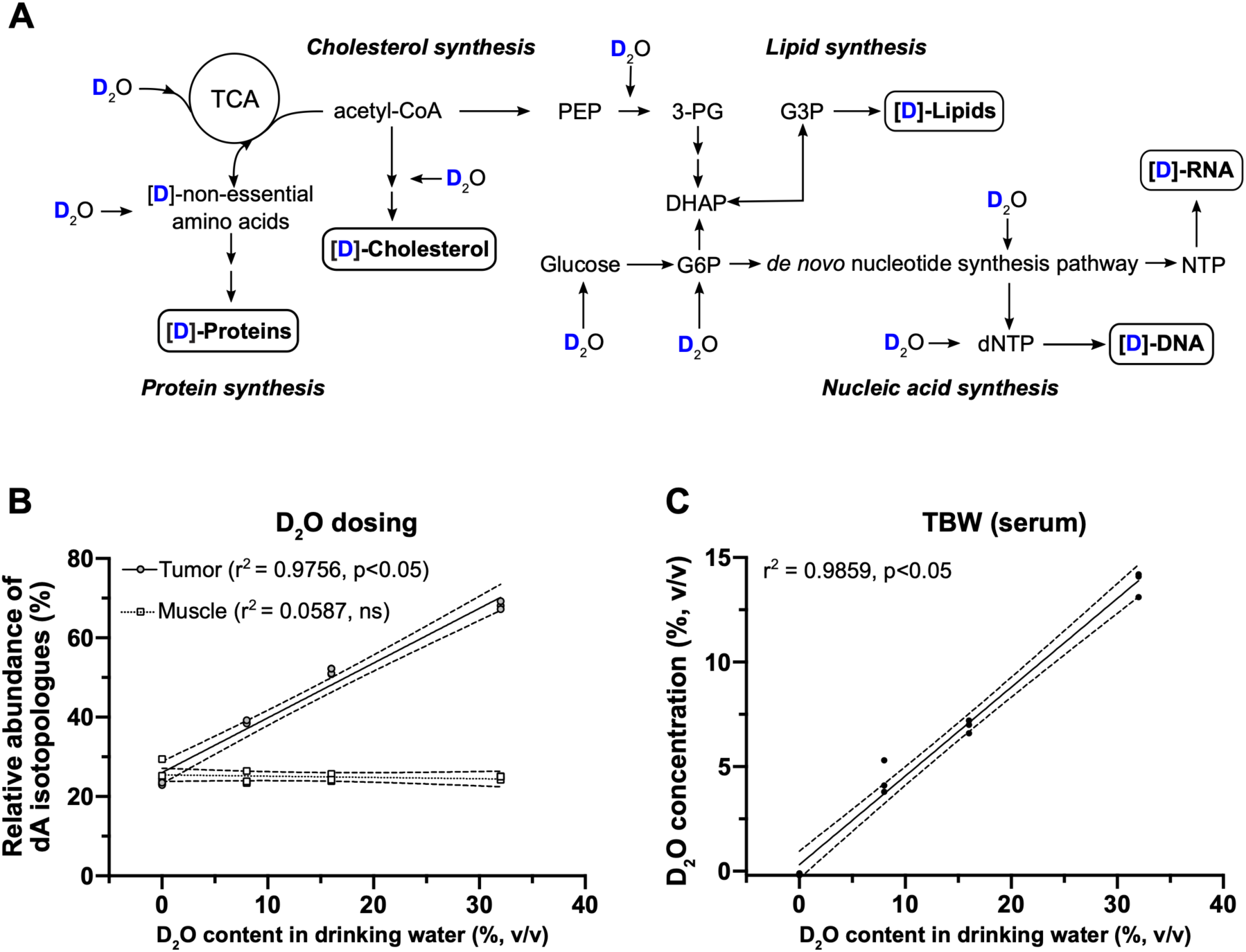
D_2_O is incorporated into biomolecules and leads to a dose-dependent increase in the isotopologue abundance of dA. **A)** Schematic illustration of biosynthetic pathways in a cell that utilize water and are therefore possible routes for nonexchangeable, stable deuterium incorporation into carbon-deuterium bonds. PEP, phosphoenolpyruvate; 3-PG, 3-phosphoglycerate; DHAP, dihydroacetone-phosphate; G6P, glucose-6-phosphate; G3P, glycerol-3-phosphate; (d)NTP, (deoxy)nucleoside triphosphate. **B)** Dose-dependent increase of dA isotopologue abundance (dA m+1, m+2, m+3, m+4 and m+5). Mice were injected with human HT-29 adenocarcinoma tumor cells (1×10^6^) in the right hind limb and labeled with increasing concentrations of D_2_O by injection of up to four i.p. boluses (35 ml/kg, 0.9% NaCl in 100% D_2_O) followed by the administration of D_2_O in drinking water for maintenance dosing (0- 32%, v/v). Two weeks after tumor cell implantation, the tumors as well as muscle tissue of the contralateral hind limb serving as a negative control were excised. The DNA was isolated and dA isotopologue abundance was quantified via GC-MS/MS. Statistical testing was performed using linear regression (n=3 per group). **C)** The D_2_O concentration in total body water was quantified using serum samples collected via mandibular or retroorbital bleed after a two-week labeling period with increasing concentrations of D_2_O. The quantification was carried out using a headspace GC-MS method and statistically tested using linear regression (n=3 per group).

**Figure 2:**
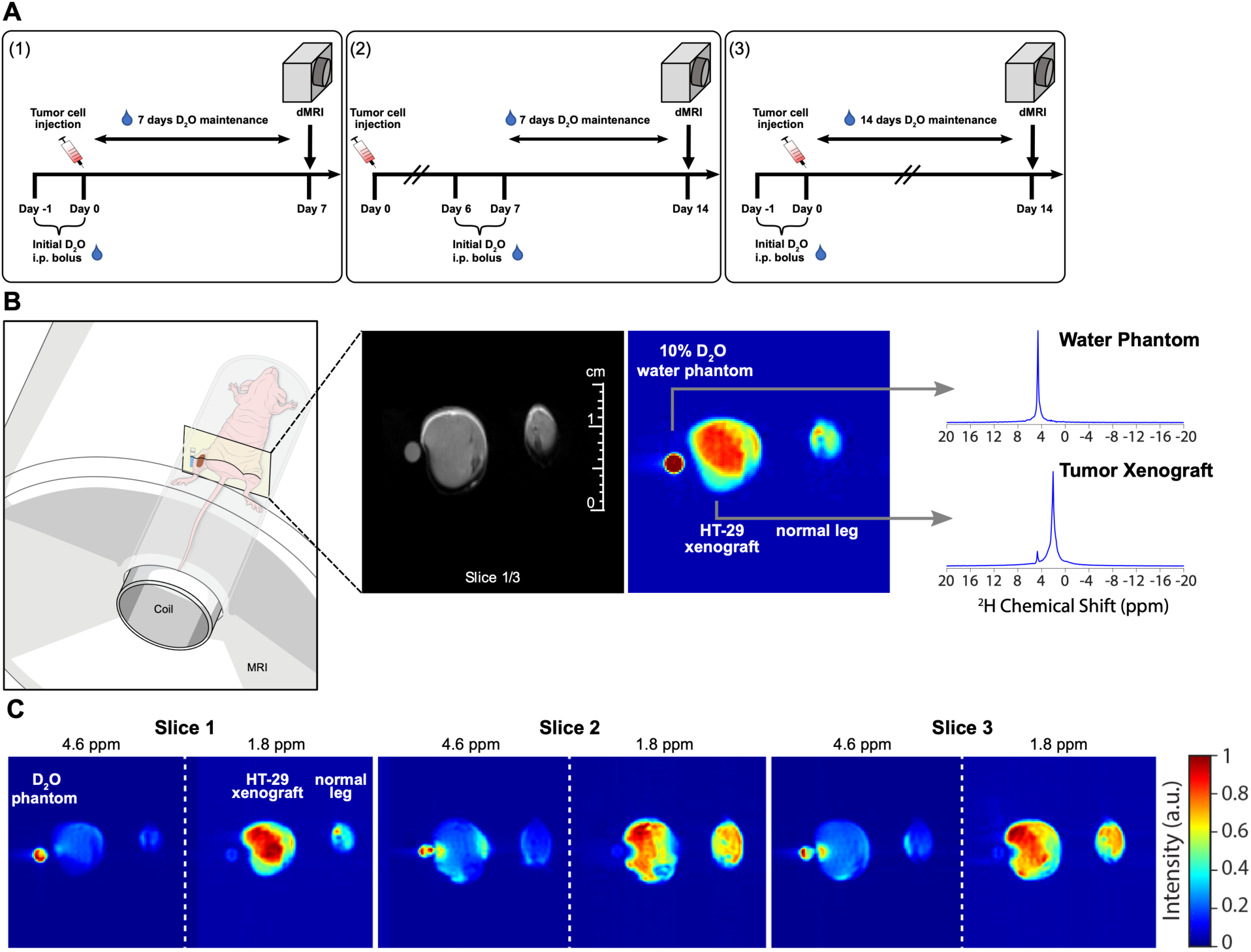
dMRSI chemical shift imaging of xenografted HT-29 tumors in mice at 11.7T. **A)** Labeling protocol for imaging experiments. Mice that underwent imaging were either labeled with two i.p. bolus injections (35 ml/kg 0.9% NaCl in 100% D_2_O) 24h apart on day −1 and 0 (1+3) or on day 6 and 7 (2) and were subsequently provided a 16% D_2_O maintenance dose in drinking water for 7 (1+2) or 14 days (3) after tumor implantation. Human HT-29 adenocarcinoma cells (1×10^6^) were subcutaneously injected on day 0 into the right hind limb of athymic Foxn1^nu^ mice. **B)** Illustration of the imaging setup (left) with a representative deuterium image (right) along with the corresponding anatomical proton MRI image (center) for labeling protocol (3). Chemical shift imaging was used to acquire a 64 × 64 grid of deuterium spectra with each spectrum corresponding to a voxel 0.5 mm × 0.5 mm × 3 mm in size. The deuterium image was formed by summing across the entire spectra while the spectra corresponding to the individual voxels indicated are shown on the right (top, D_2_O phantom; bottom, tumor xenograft). The anatomical imaging was acquired using a spin- echo sequence with TR/TE = 500 ms/ 8ms and the same geometry as chemical shift imaging but with a matrix size of 128 × 128. **C)** Spectrally selective images for the mouse from (b) corresponding to the two major peaks at 4.6 ppm (D_2_O chemical shift) and 1.8 ppm.

### Urine/serum sample collection for HS-GC-NCI analysis

Urine and blood sampling can be used interchangeably to quantify total body water D_2_O enrichment as previously shown (17). As mice recovered from anesthesia on a heating plate, ∼50 μl of urine was collected on a sheet of parafilm upon spontaneous passage. The urine was immediately transferred to a plastic microcentrifuge tube. When serum was used, it was collected via mandibular or retroorbital bleed and allowed to clot for 30 min at room temperature. The samples were spun down at 2,655 x g (5417R, Eppendorf) and the supernatant was transferred to a new microcentrifuge tube. Urine and serum samples were stored at –20 °C until TBW D_2_O enrichment analysis using headspace gas chromatography–negative chemical ionization mass spectrometry (HS-GC-NCI-MS) was performed as previously published (17).

### Tissue sample collection and preparation for GC-MS/MS enrichment analysis

To determine the isotopological enrichment of deuterium in the DNA base deoxyadenosine (dA), tissue samples from HT-29 tumors or anterior thigh muscle of the contralateral hind limb were excised immediately post-mortem after imaging and stored at −80 °C. Subsequently, we used a modified version of our validated GC-MS/MS method for analysis(18). Briefly, after isolating DNA from mouse tissue samples using a tissue DNA extraction kit (Maxwell^®^ 16, Promega), the purified DNA was incubated and hydrolyzed enzymatically (EpiQuik, Epigentek Group Inc.) to its nucleoside bases (e.g. dA, dT). The method employed solid phase extraction (Waters HLB) to extract and purify dA (unlabeled and deuterium labeled) from leg muscle and tumor tissue, with automated on-line methylation (derivatization) and rapid chromatographic analysis (∼6 min) using an Agilent GC-MS/MS system (7890A GC, LTM Series II Fast GC Module, 7000C GC-MS/MS Triple Quadrupole, 7693 Autosampler and 7697A Headspace Sampler, all Agilent Technologies). The prepared samples were injected into the GC using the following conditions (1 µL pulsed split-less injection at 235 °C; component separation using low thermal mass DB-17MS column 15 m × 0.25 mm ID × 0.25 µm film with column oven temperature program from 50-320 °C at 120 °C/min). The MS utilized positive chemical ionization (PCI with isobutane reagent gas) and full scan mode (150 to 350 Da) to acquire MS data for evaluation.

As depicted in Supplementary Fig. 1B and C, MS overlays (normalized) of methylated dA and its isotopologues (e.g. dA M+1, dA M+2, dA M+3 etc.) depict the stable isotopes of ^13^C, ^15^N, ^2^H, ^18^O found naturally (i.e. ∼23% background) in methylated dA and its isotopologues (Supplementary Fig. 1B, contralateral leg muscle, control), as well as enrichment of ∼27% deuterium (∼50% minus natural isotopic background) into the DNA base dA of rapidly proliferating cells (Supplementary Fig. 1C, HT-29 tumor).

### Proton and deuterium magnetic resonance spectroscopic imaging (dMRSI)

MRI experiments were performed on an 11.7 T (Magnex Scientific) or 9.4 T (Biospec 94/30) MRI equipped with a Bruker Avance or Avance III MRI console (Bruker-Biospin) and a custom, in-house built elliptical dual-resonance transmit/receive coil consisting of an inner elliptical solenoid deuterium coil and a saddle proton coil (Supplementary Fig. 2A). The mice were imaged with both legs perpendicular to the B0 field. The homogeneity of both coils was tested using a tight fitting 3-D printed customized oval bottle that contained two compartments with regular water and water enriched with 5% D_2_O (Supplementary Fig. 2B).

Mice were anesthetized with isoflurane (4% for induction and 1.5% – 2.5% for maintenance in medical air, 500 ml/min). During anesthesia, the respiratory rate was monitored with a pressure transducer (SA Instruments Inc.) and maintained at 60 ± 10 breaths per minute. Core body temperature was also monitored using a nonmagnetic rectal temperature probe (FISO) and maintained at 36 ± 1°C using a circulating water-warming pad. Immediately following anesthesia, both hind limbs were placed into the ^1^H/^2^H coil. A 3-mm diameter phantom tube 10% D_2_O in H_2_O (v/v) was placed adjacent to the right hind limb and was included with each scan to serve as a deuterium reference signal.

For dMRS imaging, three slices of 0.5 mm × 0.5 mm × 3 mm in size were acquired by chemical shift imaging without ^1^H decoupling using standard linear k-space encoding with a 397 ms repetition time, 512 FID points, and a sweep width of 4,000 Hz. Due to difficulties in proper phase adjustment arising from the susceptibility artifact near the D_2_O glass tube in the HT-29 images (Fig. 2 and 3), the spectra in each voxel were processed in magnitude mode. MiaPaCa- 2 MRI images (Fig. 4) were therefore acquired without a phantom vial attached to the leg. The images collected in a 64×64 matrix were then zero-filled to a final size of 128 × 128 × 3. The total scan time was 27 minutes.

**Figure 3:**
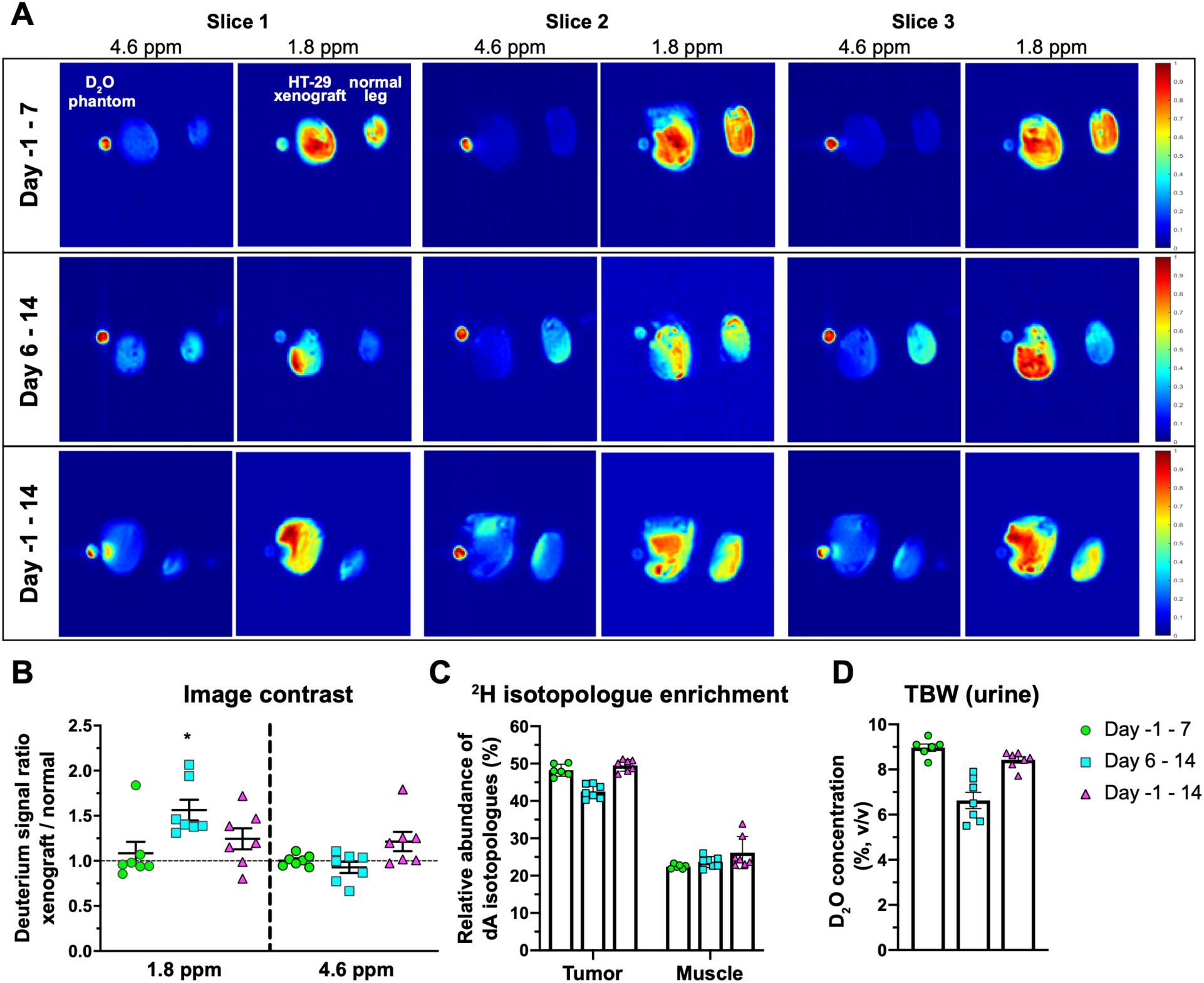
Comparison of different labeling strategies for dMRSI detection of HT-29 tumors. **A)** Representative images of the three labeling protocols shown as spectrally separated images for the 1.8 ppm and 4.8 ppm (water) peaks across three slices of HT-29 tumors acquired on a 11.7T scanner. **B)** Quantification of the contrast between tumor and control hind legs for the labeling schemas defined in Fig. 2a summed across the leg volume for the aliphatic (1.8 ppm) and water (4.6 ppm) signal regions. The leg area was defined as an ROI, the mean grey value was measured within and expressed as a ratio between the HT-29-injected and normal leg. Statistical analysis was performed using a Wilcoxon signed rank test against a hypothetical median of one, *=p<0.05, n=7 per group. **C)** Relative abundance of dA isotopologues after a 7-day or 14-day labeling period in tumor and muscle tissue (n=6-7 per group) analyzed via GC-MS. **D)** The D_2_O concentration in total body water was quantified using urine samples collected after 7 and 14 days of labeling via HS-GC-MS, n=6-7 per group.

**Figure 4:**
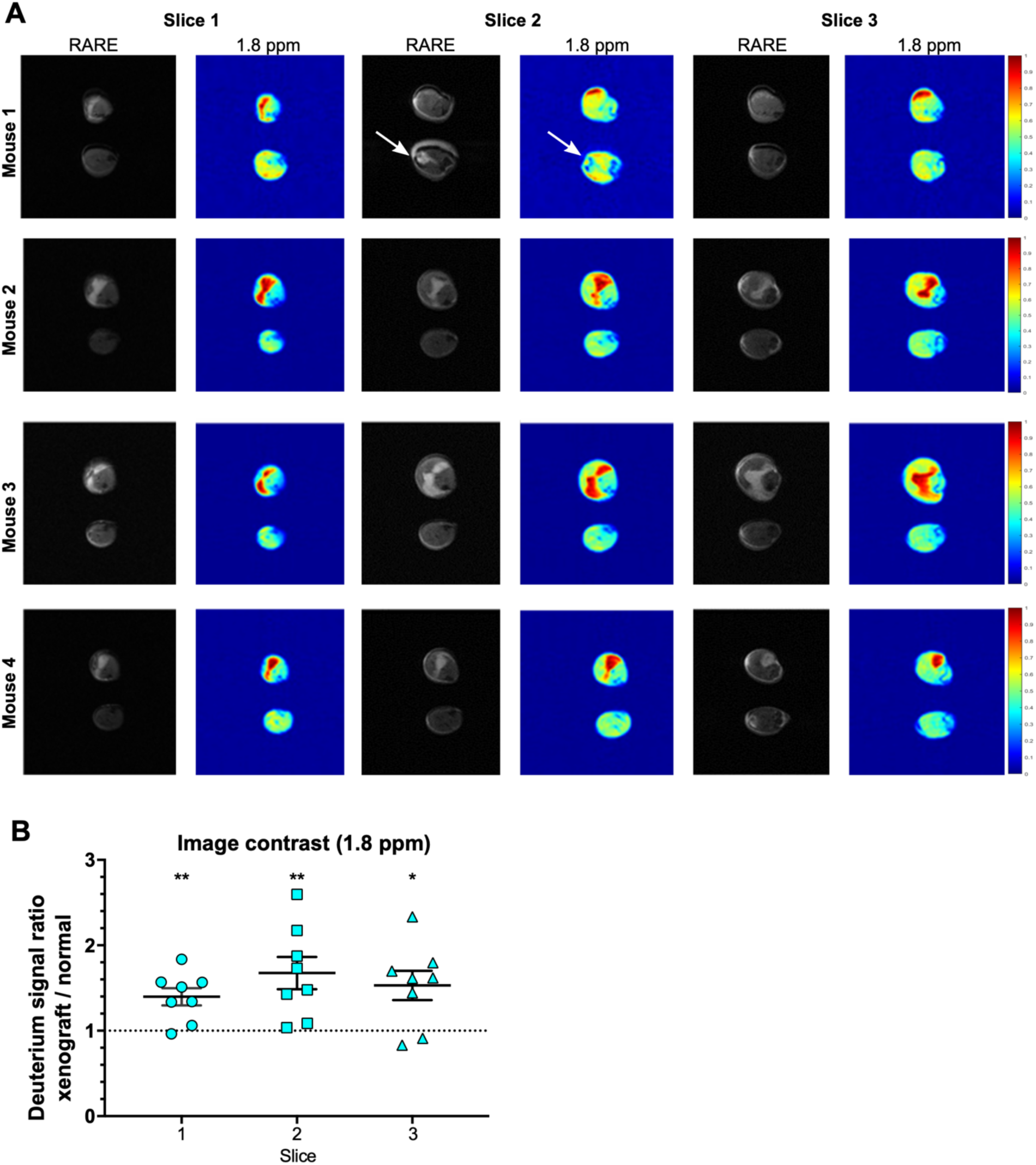
Detection of MiaPaCa-2 tumors via dMRSI. **A)** Paired RARE anatomical and deuterium images of 4 representative mice at 1.8 ppm of the MiaPaCa-2 xenograft-injected leg (top) and the contralateral side (bottom) after two weeks of tumor growth and D_2_O labeling during the second week. The white arrows indicate a hyperintense tendon present on the RARE MR image that is not detectable in the corresponding deuterium image. Images were acquired using a 9.4T scanner. **B)** Quantification of the contrast between tumor-bearing and control hind legs for the labeling schemas 2 as defined in Fig. 2A summed across the leg volume for the aliphatic (1.8 ppm) signal region for each of the three acquired images per mouse. Statistical analysis was performed using a one sample test against a hypothetical median of one, n=8, *=p<0.05, **=p<.0.01. Error bars indicate the SEM.

To increase the sensitivity to the weak background D_2_O signal, a noise reduction algorithm was employed. It takes advantage of the repeating structure of the data by decomposing the 4D image into a multilinear combination of vectors representing spectra and image columns and rows by low rank Tucker Decomposition. The algorithm uses 32 spectral vectors, 32 image vectors in each direction and all 3 slice vectors to reconstruct the original the 512 × 128 × 128 × 3 image (19). Using this method, 92 ± 4% of the variance was captured in each scan, with the residual approximately corresponding to the noise level by visual analysis. Noise reduction was primarily required to detect the weaker D_2_O signal at 4.6 ppm - maps of the stronger metabolite signal at 1.8 ppm were similar in the presence and absence of noise reduction. The water peak from HDO was set to 4.6 ppm relative to the DSS value, following a temperature correction to 37 °C (20). No signal was detected when D_2_O labeling was omitted (Supplementary Fig. 2C). The anatomical imaging was performed using the following parameters: spin-echo sequence (RARE, rapid acquisition with refocused echoes), TR/TE = 500 ms/ 8ms, with the same geometry as chemical shift imaging but with a matrix size of 128 x 128.

To compare metabolism between tumor and non-tumor areas, a spectrally selective image was first formed by summing the 10 points (156 Hz) on either side of the two major peaks in the spectra after noise reduction. A region of interest was then drawn around the right and left leg for each of the three slices in ImageJ (21). The mean greyscale value was tabulated for each region and a ratio between the signal intensity of the tumor-injected and the normal leg was calculated for both spectrally selective images.

### *Ex vivo* NMR

Metabolites from tumor sections frozen at −80 °C were subjected to three-phase separation to produce a polar phase, an insoluble protein interphase and a non-polar phase containing lipids using modifications to a previously published procedure for cell extracts (22,23). The non-polar fraction was prepared for deuterium NMR by dissolving the lyophilized powder in 75% chloroform/25% methanol, with 5% deuterated chloroform present for the frequency lock. Samples for proton NMR were prepared in deuterated 75% chloroform/25% methanol. Polar samples were dissolved in 90% H_2_O/10% D_2_O. Spectra were recorded at 16.4 T at 20°C in a Bruker spectrometer equipped with a 3 mm triple inverse resonance cryoprobe. 512 transients were acquired for each sample. A presaturation sequence was used for the proton spectra with a relaxation delay of 6 seconds, a pulse acquire sequence was used for the deuterium spectra with a relaxation delay of 2 seconds.

### Statistical analysis and software

For HS-GC-MS/MS operation, data acquisition and processing, we utilized a PC workstation with Agilent MassHunter Acquisition (B.07.06.2704), Quantitative (B.08.00) and Qualitative (B.07.00, Service Pack 1) software. Data were analyzed and visualized using GraphPad Prism 8.0. All graphs represent the mean value ± SEM. Statistical analysis was performed as indicated in the figure legends with p<0.05 considered significant.

## Results

### D_2_O uptake leads to a dose-dependent increase of isotopologically labeled deoxyadenosine in tumor cells

Water is a substrate for multiple biosynthetic pathways required for cell proliferation, including amino acid (proteins, peptides), nucleic acid (DNA, RNA), and fatty acid (lipids) synthesis (Fig. 1A). When deuterated water is administered systemically, biomolecules within rapidly proliferating cells incorporate deuterium; hence, deuterium enrichment can serve as a proxy for cell proliferation (8,9,13,24-26). Using a quantitative gas chromatography tandem mass spectrometry (GC-MS/MS) method, we measured deuterium enrichment of deoxyadenosine (dA) in xenografted tumors following 7 and 14 days of *in vivo* tumor growth during concurrent D_2_O labeling (13,17,18). The contralateral quadriceps muscle was excised to serve as control tissue for the GC-MS/MS analysis. In addition, we collected urine and serum samples from each study animal on the day of imaging to measure total body water (TBW) deuterium enrichment via headspace (HS)-GC-MS (17), which was confirmed to be ∼6-9% for all animals that underwent imaging.

The natural isotopic background in deoxyadenosine, resulting mainly from the natural abundance of ^2^H, ^13^C, ^15^N and ^17^O, ranges from 23% to 26% in both tumor without D_2_O labeling and muscle tissue (Fig. 1B). Deoxyadenosine extracted and purified from the tumors of mice receiving maintenance deuterated water of increasing concentrations (0%, 8%, 16% and 32%) showed a dose-dependent linear increase in isotopologue enrichment of dA.

This increase in dA isotopologue enrichment above natural isotopic background was not detected in muscle tissue of the contralateral leg (Fig. 1B). Metabolic labeling of cellular biomass with 8% D_2_O as maintenance dose (in drinking water) resulted in ∼5% D_2_O in TBW and a tumor deuterium enrichment of 38.7% ± 0.2 after two weeks of labeling (Fig. 1B, C). Higher maintenance doses of D_2_O led to an increased TBW and tumor dA enrichment, i.e. 16% D_2_O resulted in ∼7% TBW and 51.3% ± 0.4 dA isotopologue enrichment and 32% D_2_O resulted in ∼14% TBW and 68.2% ± 0.6 dA isotopologue enrichment, respectively (Fig. 1B, C). Meanwhile, the isotopologue enrichment of dA isolated from the contralateral muscle tissue (control) remained stable at around 24% for the full range of tested TBW D_2_O concentrations (Fig. 1B), confirming that D_2_O preferentially labeled tumor cells. Additionally, we found that the use of 8%, 16%, and 32% D_2_O in drinking water led to multiple deuterium atoms being incorporated into the dA molecule, with more than half of dA molecules containing more than one deuterium atom (e.g. dA M+2, M+3, Supplementary Fig. 1A). Although we can reliably quantify the deuterium enrichment via mass spectrometry, the analysis is carried out *ex vivo* and requires a tissue sample. In a clinical setting, acquiring tissue biopsies can be difficult and there are situations where the tumor is inaccessible, or the biopsy is otherwise medically inadvisable.

### Deuterium incorporated into biomolecules during D_2_O labeling is detectable by dMRSI

In addition to clinical contraindications for biopsy, the heterogeneity and variability in metabolic activity within a tumor may result in inconsistent deuterium quantification depending on the specific region sampled (27,28). Due to these limitations, we pursued the development of an imaging modality that would circumvent the clinical risks and potential sampling error of the biopsy approach. We hypothesized that *in vivo* imaging of deuterium following biosynthetic labeling would allow noninvasive detection and monitoring of molecular pathways distinct from those visualized with current techniques such as FDG-PET or hyperpolarized ^13^C-MRI. DNA itself is unlikely to be detected via MRI due to its short T2 and long T1 relaxation rates stemming from its size and relative stiffness. However, other metabolic intermediates enriched with deuterium may be detected, if they accumulate within the tumor in sufficient abundance (29,30). We therefore set out to evaluate whether higher deuterium content within tumors compared to healthy normal tissue, as a result of increased biosynthetic rates of the former, would allow distinct visualization of tumors via dMRSI.

We performed *in vivo* labeling on mice injected with HT-29 tumors followed by dMRSI detection (Fig. 2A). Figure 2B shows a representative slice from the image of both hind limbs of a female athymic mouse with the left leg bearing a relatively small (<1 cm) HT-29 tumor xenograft taken 14 days after tumor cell injection with concurrent deuterated water labeling to 8% TBW. A 3-mm NRM capillary tube containing 10% D_2_O near the left leg served as a reference for the frequency of the water signal at ∼4.6 ppm. Two major peaks can be detected in the deuterium spectra. The first is centered at 4.6 ppm near the expected frequency of the HDO/D_2_O water peak (top). There is a close correspondence between the anatomical MRI and the deuterium image at the water frequency, indicating background TBW deuterium enrichment is similar between the tumor-bearing limb and the unmanipulated limb (Fig. 2C). The remaining, non-water derived signal is characterized by an intense peak (mean SNR ∼150) with a chemical shift of ∼1.8 ppm (Fig. 2B, bottom spectrum) that is primarily, but not exclusively, observed in the tumor region. Figure 2C shows an example of all three slices acquired and separated into the two distinct frequencies, demonstrating that the tumor deuterium signal is not attributable to free D_2_O. In all three slices, the deuterium signal intensity at 1.8 ppm is highest in the tumor bearing limb. The exact identity of the peak cannot be directly inferred from MRI, but the shift indicates that the peak likely originates from the resonance of alkyl groups, taking into account the slight chemical shift offset from B_0_ distortion caused by proximity to the partially filled D_2_O phantom vial (31).

### dMRSI distinguishes tumors from normal tissue

To detect deuterium-labeled metabolites *in vivo*, we tested several labeling-imaging protocols. Throughout the tested experimental schemas, we consistently observed a stronger deuterium signal at ∼1.8 ppm in the tumor region compared to the contralateral limb (Fig. 3A). The degree of contrast, however, was dependent on both the length and timing of the labeling period (Fig. 3B). The strongest contrast (median=1.40, p= 0.02 on a per mouse basis, Wilcox’s signed rank test), quantified as a ratio of the mean grey value between the HT-29-injected and normal leg, was observed after 14 days of tumor growth with labeling starting on the 6^th^ day (labeling schema (2) in Fig. 2A, cyan squares in Figure 3B). Limiting both growth and labeling to the first 7 days (labeling schema (1) in Fig. 2A) led to negligible contrast (median=0.96, p=0.81). A similar trend was observed in an independent experiment performed on a 9.4 T MRI (Supplementary Fig. 3). Labeling for the full 14-day tumor growth period (labeling schema (3) in Fig. 2A) decreased contrast relative to starting at the 6-day midpoint (median=1.20, p=0.11). No significant difference in contrast was observed comparing the signal intensity at 4.6 ppm for any labeling protocol. Interestingly, while the MRI contrast was strongly dependent on the labeling period, the degree of deuterium labeling in both dA and in TBW was nearly independent of the labeling period (Fig. 3C, D).

To validate our findings, we tested the most promising labeling strategy (d 6 – 14) in a second xenograft model using the pancreatic cancer cell line MiaPaCA-2 (Fig. 4). Tumor cells were again subcutaneously injected into the right hind leg and imaged after 14 days. As previously demonstrated with the HT-29 cell line, deuterium labeling from day 6 to 14 provided a significant contrast, enabling a clear distinction between the tumor-injected leg and the contralateral limb (Fig. 4A, B). In contrast to HT-29 tumors, MiaPaCa-2 tumors are easily identifiable on the anatomical MRI and the comparison of the deuterium signal with the anatomical image confirmed that the highest deuterium signal originates in the tumor tissue. Of note, other hyperintense anatomical structures such as tendons or the subcutaneous fat were not detectable in the deuterium image (Fig 4A, white arrows).

### dMRSI signal from D_2_O labeling originates from cholesterol

The deuterium signal at ∼1.8 ppm likely arises from alkyl groups, but due to the spectral overlap in this region and the limited *in vivo* resolution the peak cannot be assigned to a specific metabolite. To more definitively assign the peak, tumors were excised, and flash frozen at the end of the labeling period immediately following imaging. The polar and non-polar fractions were separated and analyzed by deuterium NMR (Fig. 5). No significant peaks were detected in the deuterium spectrum of the polar fraction beyond the HDO solvent signal (data not shown), suggesting TCA cycle intermediates and amino acids are rapidly cycled and do not accumulate to a substantial degree within the experimental timeframe. Within the non-polar fraction, no signal outside of those of the solvent could be detected in an unlabeled tumor or for the day −1-7 labeling period (Fig. 5B-D), in concordance with the weak signal observed by *in vivo* dMRSI. When labeling occurs during the period of rapid tumor growth, two peaks at 0.7 ppm and 1.1 ppm could be detected with D_2_O labeling for both HT-29 (Fig. 5F, H) and MiaPaCa-2 xenografts (Fig. 5E, G). The peak at 0.7 ppm occurs in an empty region of the corresponding proton spectra and as such can be immediately identified as the distinctive high intensity singlet peak of the methyl group of carbon 18 of cholesterol and cholesterol esters based on previously assigned spectra (32,33). The peak at 1.1 ppm occurs in a somewhat more crowded region and is less distinctive but is consistent with the corresponding C19 methyl group. Only a minor peak for the cholesterol ester was detected in a muscle sample from the contralateral limb (Fig. 5I). No peaks for lipids or other non-polar molecules in either the xenograft or muscle samples were detected in the deuterium NMR, despite their presence in the proton spectra (Fig. 5A).

**Figure 5:**
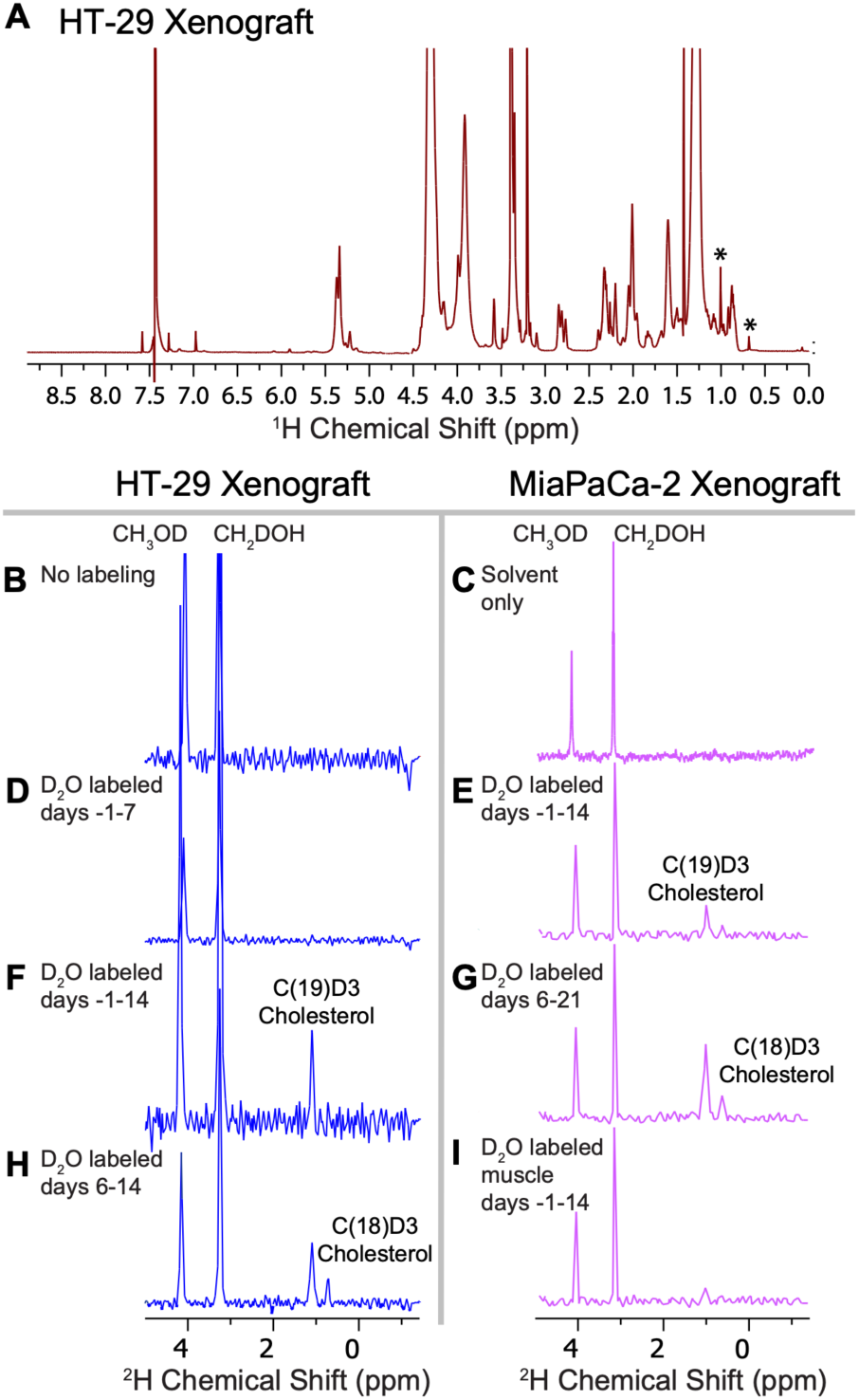
NMR spectra of the non-polar fraction of extracted xenografts. **A)** Proton NMR spectrum of the non-polar fraction from a HT-29 xenograft. Numerous peaks corresponding to lipids are visible as well as two peaks corresponding to the strong singlet signal of C(18)D3 and C(19)D3 of cholesterol and cholesterol esters (marked with *) at 0.7 ppm and 1.1 ppm. **B and C)** Deuterium NMR spectrum of the non-polar fraction from a HT-29 xenograft without D_2_O labeling. No peaks are present beyond those of the solvent. **D to H)** Non-polar fractions in the presence of D_2_O labeling. Peaks at the expected position for C(18)D3 and C(19)D3 of cholesterol and cholesterol esters are evident in both xenografts when D_2_O is administered during the tumor growth period **(E-H). I)** Deuterium NMR spectrum of the non-polar fraction from muscle tissue from the contralateral limb. Only a weak signal from labeled cholesterol esters is evident. Spectra are referenced to the CDCl_3_ peak at 7.23 ppm. Intensities are normalized to the dry protein weight of the sample. No peaks of any type were found in the corresponding polar fractions.

## Discussion

The first published use of deuterium in MRI was described in 1986 (34). Initial studies on dMRS showed that although deuterium has a natural abundance of only 0.015% and a gyromagnetic ratio that is 1/6^th^ of a proton, other properties, such as its short T1 relaxation time that allows for quicker signal averaging, can compensate for the less favorable MRI-relevant factors (34). Deuterium MRI has since been used to visualize blood flow within a variety of tissues in animal models, including tumors (35-38). In contrast to our approach, most studies of this type used a single bolus injection of deuterated water to trace the deuterium signal in the organ of interest over a short time frame of minutes to hours. This enabled the visualization of blood flow and vasculature allowing the evaluation of tumor perfusion, but perfusion alone does not discriminate between healthy tissue and tumor. Continuous labeling with deuterium rather than administering a bolus of deuterium-labeled metabolite enabled us to enrich the tumor tissue with deuterium through biosynthesis of molecules incorporating deuterium. Unlike deuterium imaging following a single bolus injection before image acquisition, the D_2_O in our protocol had equilibrated with the TBW; i.e. we were not evaluating a perfusion difference but rather the accumulation of recently synthesized biomolecules within the tumor. We have used dA deuterium enrichment to demonstrate and quantify the degree of nucleotide labeling, which provides an estimate of the total degree of biosynthetic labeling with deuterium (17,18) as DNA is rarely catabolized once synthesized (39). Deuterium has also been shown to be incorporated into other major classes of biomolecules, such as amino acids/proteins (9,24), triglycerides/lipids (40-42), glycolysis (43) and TCA cycle intermediates (44). Therefore, we inferred that many biomolecules are labeled with deuterium during the experiment, and likely at multiple sites (see Supplementary Fig. 1B). However, the labeled molecules are also subject to excretion and conversion into other metabolites as time progresses. Furthermore, the metabolism of tumors is not closed but contains substantial contributions from circulating molecules synthesized elsewhere. The localized deuterium signal of a particular metabolite is therefore a measure of the net accumulation at the site from all internal and external sources from molecules synthesized within the specified timeframe.

By testing different labeling schemas, we found that labeling from day 6 – 14 provided the highest contrast between the tumor-injected leg and the contralateral control. During this time period, the tumor doubles in size resulting in high deuterium enrichment and thus a readily detectable tumor (16). More prolonged labeling with D_2_O reduced the contrast between the proliferating tumor and muscle in the contralateral limb. Although the muscle tissue showed no significant increase in isotopologue enrichment of dA after two weeks of labeling, other biomolecules that contribute to the imaging but are not readily turned over, i.e. lipids and sterols, may enrich with deuterium over time. This was previously observed with long term D_2_O labeling of healthy tissues (45,46). The labeled metabolite generated will depend on the specifics of the metabolism of the system (42,47).

Within the selected labeling window, we observed accumulation of cholesterol and cholesterol esters in HT-29 (colon adenocarcinoma) and MiaPaCa-2 (pancreatic ductal adenocarcinoma) xenografts (Figs. 4 and 5). Cellular cholesterol is obtained either from circulating LDL-cholesterol in plasma by receptor-mediated uptake or by *de novo* synthesis within the tumor via the mevalonate pathway with acyl-CoA as the carbon source. The degree of labeling is dependent on the source. Cholesterol obtained directly from dietary sources is exogenous to the body, not subject to D_2_O labeling and, therefore, invisible to dMRSI. Cholesterol formed from other precursors, whether obtained by receptor mediated endocytosis of circulating cholesterol or by on-site synthesis, is likely to be heavily labeled by D_2_O (48). While circulating cholesterol from dietary sources forms a large fraction of total cholesterol in most normal tissues, *de novo* synthesis of cholesterol is strongly upregulated in many tumors (49) from proteolytic cleavage of sterol regulatory element binding proteins (SREBPs) in response to p53, RAS, or mTOR activation (50) originating from oncogenic mutations or hypoxic conditions and other environmental stressors. Concurrently, cholesterol efflux is often reduced via downregulation of the ATP-binding cassette transporter A1 (ABCA1) through the effects p53 and RAS on DNA hypermethylation (51). The result is a net accumulation of cholesterol in the membranes and cholesterol esters in lipid droplets. Membrane cholesterol is likely not detectable in MRI due to excessive signal broadening from restricted motional averaging within the bilayer. Cholesterol esters in lipid droplets, by contrast, are relatively mobile and are expected to give sharp MRI signals (31). Surprisingly, we saw little evidence of labeled lipids (Fig. 5), another major potential synthetic route for long term storage.

Cholesterol accumulation within the tumor has many consequences (50). High levels of cholesterol in the mitochondrial membrane alter its physical properties, inhibiting the mitochondrial permeability transition as well as the activation of apoptosis by limiting the release of cytochrome C (52). Cholesterol binding directly activates mTORC1 (53), the Smoothened receptor (54), and multiple signaling proteins containing PDZ domains with the CRAC motif (55) resulting in the activation of several major pathways associated with cell division, differentiation, invasion, and metastasis including the MAPK/ERK, Hedgehog, PI3K/Akt, and Wnt/β-catenin pathways (50). Indirectly, high levels of membrane cholesterol can facilitate the association of aberrant signaling complexes through the excessive formation of lipid rafts (56). Given the multiple oncogenic pathways cholesterol affects, it is not surprising that cholesterol levels are often strong predictors of cancer progression (57) and that modulating cholesterol metabolism is an active target for drug development.

Since cholesterol can be derived from many sources, drug development targeting cholesterol metabolism is contingent on reliable methods for the measurement of cholesterol uptake, synthesis, and accumulation. The primary method for cholesterol imaging is currently the SPECT tracer NP-59, an ^131^I cholesterol analogue (58). As a SPECT tracer, NP-59 measures levels of cholesterol uptake, but not *de novo* synthesis, which is an important part of cholesterol metabolism in many tumors (49). By imaging accumulation of synthesized cholesterol, deuterium imaging offers a window into an important aspect of tumor metabolism that has been difficult to access by imaging techniques to date.

From a clinical translation standpoint, deuterium MRI is inexpensive with regard to label and system set-up, technically straightforward regarding the design of RF coils for lower frequencies, and it can be used to evaluate multiple anatomical regions. Short-term intake of low-to-moderate concentrations of deuterium is generally considered safe for humans and long-term toxicity studies in animals have not shown any detrimental effects below 20% TBW enrichment (11). Nonetheless, some logistical considerations must be overcome for clinical implementation of this imaging approach. Clinical studies involving deuterated water administration have so far been performed at 1-2% TBW enrichment (59,60). Future pre-clinical studies will test lower TBW doses of D_2_O labeling for this imaging approach. Given the abundance of signal observed at the current TBW labeling it is likely that the lower doses will allow imaging resolution between tumor and healthy tissue. Once the lowest systemic pre- clinical labeling dose is determined, pilot clinical studies will be performed to determine safety. In addition, current experiments were conducted at a high field (9.4 and 11.7 T). Given the strength of the observed dMRS signal in our preclinical study (overall mean SNR∼150) and the signal gain achieved by post-acquisition processing (∼24 fold), testing our labeling/imaging protocol at lower TBW enrichments and lower magnet field strengths that are widely available for clinical imaging (i.e. 3 T) should be feasible. Furthermore, limb tumor xenografts were used to demonstrate the potential of the method, while future studies of dMRS imaging in orthotopic tumor models will be carried out to assess image contrast in a physiological context of complex anatomy.

In summary, we have demonstrated that D_2_O labeling leads to differential incorporation of deuterium into cholesterol in two distinct xenograft models that can be visualized by dMRSI *in vivo*. We believe that this novel imaging technique could provide an excellent addition to existing cancer imaging approaches and facilitate the analysis of tumor cell cholesterol metabolism *in vivo*.

## Acknowledgements

We thank Dr. Nobu Oshima for technical assistance with *in vivo* injection of HT-29 cells and Dr. Ehydel Castro for performing phlebotomy on a subset of study animals. We acknowledge Drs. Steve J. Dodd, Martin J. Lizak, Kazutoshi Yamamoto and Danielle Donahue for assisting with MRI scan acquisition. We thank Christopher Johns for assistance with tissue processing and editing of the manuscript. We acknowledge Dr. Charles Zhu for assistance with 3D MRI coil design. We thank Ethan Tyler and Erena He, medical illustrators with NIH Medical Arts, for generating the drawing used in Figure 2. This was work was supported by NCI, NIH intramural funding including a supplement from the NCI Center for Cancer Research Major Opportunity: The Metabolic Basis of Cancer.

## Competing interests

NPB, DEF, NM, HM, MCK, and REG are inventors on a patent application related to this work, PCT/US2017/058886. Additionally, JRB, SK, HM, MCK are inventors on patent application pertaining to the signal-to-noise reduction algorithm, PCT/US2018/018217.

## Notes

### Summary of Updates

Method verified in second tumor xenograft model (Fig. 4) and signal origin identified via ex vivo NMR of tumor extracts (Fig. 5). Results and Discussion section updated to reflect new findings.

