## Supplemental Figures 1-3 for "*In vivo* deuterated water labeling allows tumor visualization via deuterium magnetic resonance spectroscopic imaging of cholesterol"

**Supplementary Figures**

Julian C. Assmann^1†^, Jeffrey R. Brender^2†^, Don E. Farthing^1^, Keita Saito^2^, Shun Kishimoto^2^, Kathrynne A. Warrick^1^, Natella Maglakelidze^1^, Daniel R. Crooks^3^, Hellmut Merkle^4^, Ronald E. Gress^1^, Murali C. Krishna^2^, Nataliya P. Buxbaum^1*^

^1^Experimental Transplantation and Immunotherapy Branch, National Cancer Institute, National Institutes of Health, Bethesda, Maryland, USA

^2^Radiation Biology Branch, Center for Cancer Research, National Cancer Institute, National Institutes of Health, Bethesda, Maryland, USA

^3^Urological Oncology Branch, Center for Cancer Research, National Cancer Institute, National Institutes of Health, Bethesda, Maryland, USA

^4^Laboratory for Functional and Molecular Imaging, National Institute of Neurological Disorders and Stroke, National Institutes of Health, Bethesda, Maryland, USA

†contributed equally

^*^Corresponding author:

Nataliya P. Buxbaum, M.D., Experimental Transplantation and Immunotherapy Branch, National Cancer Institute, Bethesda, Maryland, USA,, Phone: (240) 760-6157

**Supplementary Figures**

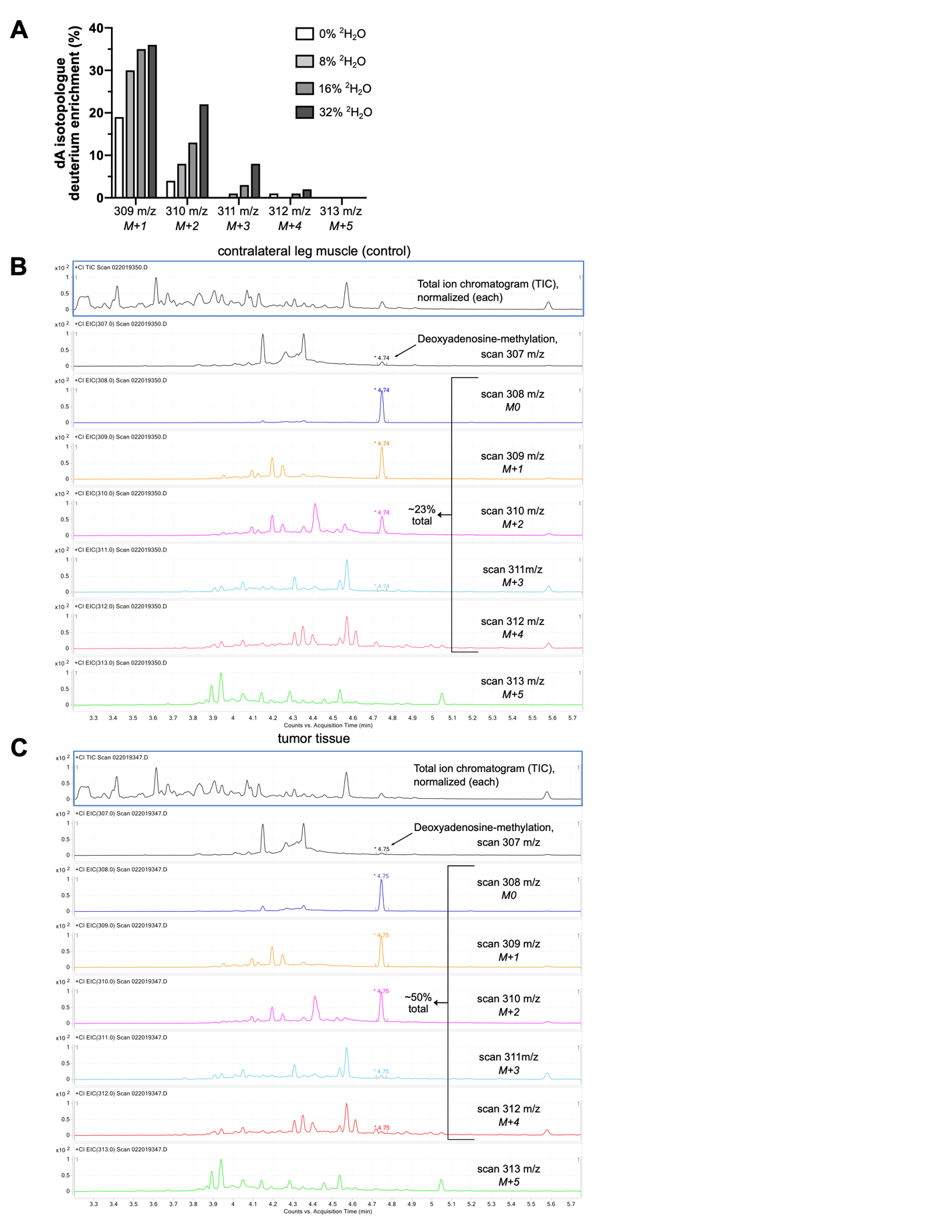

***Supplementary Figure 1: Representative result of the GC-MS analysis to quantify the abundance of dA isotopologues. a)*** *Samples of 8%, 16% and 32% ^2^H_2_O in H­_2_O from drinking bottles (one sample each) were collected at the end of the experiment. The concentration of ^2^H_2_O in drinking water was quantified via headspace GC-MS.* ***b)*** *Representative plot of four samples taken from Fig. 1a with increasing amounts of ^2^H­_2_O labeling and their respective isotopic signatures* ***c)*** *Overlay of normalized MS scans (308, 309, 310, 311, 312, and 313 m/z) representing the DNA base pair (dA) extracted and purified from a mouse leg muscle and a xenografted HT-29 tumor. The mouse was dosed to ~8% ^2^H_2_O in TBW (v/v) for this experiment. Summation of the relative abundances of the methylated dA component (retention time 4.74 min) for each dA isotopologue (e.g. dA M+1 (309 m/z), dA M+2 (310 m/z) etc.) in the muscle shows ~23% natural isotopic background (e.g. ^13^C, ^15^N, ^2^H, ^18^O) from the methylated dA component while the tumor tissue shows enrichment of ~27% deuterium (~50% minus ~23% natural isotopic background) into the DNA base pair (dA) of the HT-29 tumor cells.*

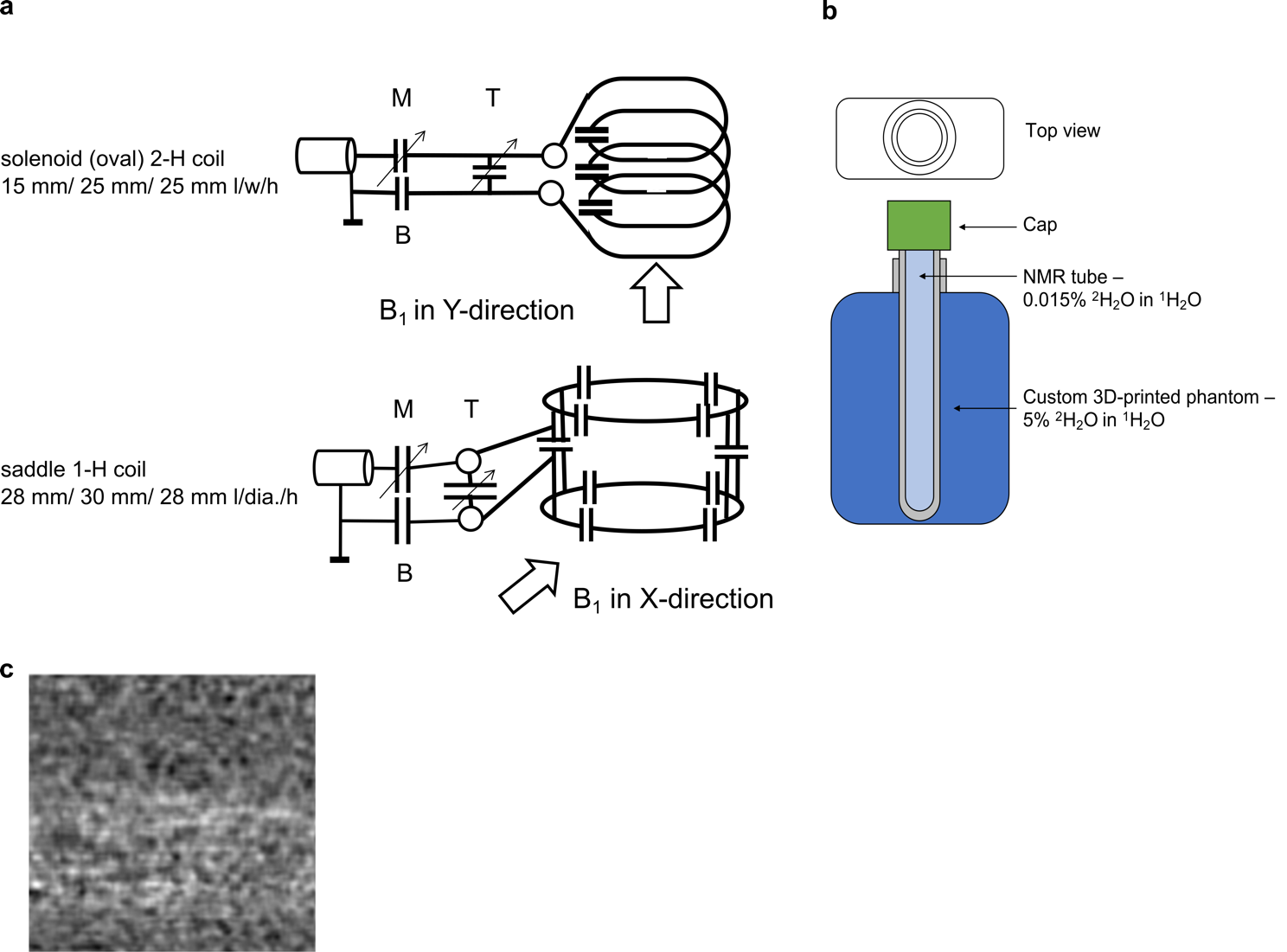

***Supplementary Figure 2: MRI coil and phantom schematics. a)*** *^1^H/^2^H transmit-receive coil schematic of the RF ^1^H saddle and oval ^2^H coil used in this study. The probes (coils) are used in transmit/receive mode within the MR system. The ^2^H coil wires (silver-plated 1.6 mm thick varnished copper) were tightly wound around the oval former and contain several in-line capacitors. The wires were attached to a standard tune/match circuit that transforms its impedance to the 50 Ω characteristic impedance of the RF transmission line. The wires of the ^1^H coil were wrapped around the shell of the solenoidal coil at a distance and were independently mechanically supported. This saddle coil had two saddle-shaped branches that were connected in parallel to a standard tune/match circuit for 50 Ω characteristic impedance RF transmission line. Both branches had in-line capacitors and branch coupling capacitors to null undesired interaction of the two parallel branches.* ***b)*** *Schematic of the custom-built phantom vial used in (c). The phantom vial is filled with regular water (0.015% natural abundance of ^2^H) on the inside and 5% ^2^H_2_O in regular water in the outer compartment.* ***c)*** *Example of a processed MR image taken from a mouse leg without labeling or tumor.*

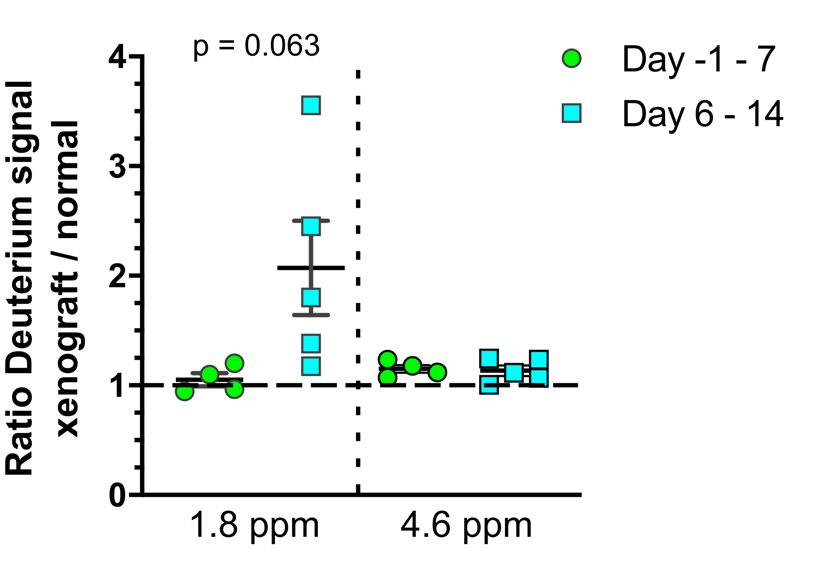

***Supplementary Figure 3: Quantification of the deuterium signal contrast ratio in xenografted HT-29 tumors at 9.4T.*** *Quantification of the contrast between tumor and control hind legs for the labeling from day -1 – 7 or 6 – 14, summed across the leg volume for the aliphatic (1.8 ppm) and water (4.6 ppm) signal regions. The leg area was defined as a ROI, the mean grey value was measured within and expressed as a ratio between the HT-29-injected and normal leg. Statistical analysis was performed using a Wilcoxon signed rank test against a hypothetical median of one, *=p<0.05, n=4-5 per group.*
